# On rRNA gene cluster variation, phylogeny and local ecological differentiation in the *Jaera albifrons* complex (Crustacea: Isopoda)

**DOI:** 10.64898/2025.12.31.697188

**Authors:** Andrey Rozenberg, Vadim Khaitov

## Abstract

The *Jaera albifrons* species complex comprises several closely related species of intertidal marine isopods that exhibit limited morphological differentiation and varying degrees of hybridization. In this pilot study, we analyzed regions of the nuclear rRNA genes (D1-D2 and ITS1) from populations of *J. albifrons*, *J. praehirsuta*, and *J. ischiosetosa* to assess their utility as diagnostic markers for species identification and phylogenetic inference. Despite the species’ morphological distinctness and ecological differences, rRNA variability was remarkably shallow across the complex, with only few variable sites in D1-D2 and ITS1. This contrasted sharply with high interspecific divergence observed in other *Jaera* species, indicating recent divergence and nascent speciation within the complex. *J. ischiosetosa* showed the lowest intra-individual variation and was most distinct from the two other species. Overlap between *J. albifrons* and *J. praehirsuta* genotypes, consistent with known introgressive hybridization, was considerable and likely stems from introgressive hybridization. Local hybridization patterns identifiable morphologically varied across settlements with of the settlements demonstrating unusually high hybridization rates. Different settlements were found to demonstrate a range of abiotic factors impacting spatial separation of the different species and varying degrees of similarity between the species in rRNA genotypes. Phylogenetic analysis of the genus revealed that the *J. albifrons* complex originates from among the species of the Mediterranean species group and represents a recent migration from the Mediterranean rather than an early-diverging lineage as previously suggested. These findings highlight the challenges of using rRNA markers for species delimitation in recently diverged and hybridizing taxa and underscore variable hybridization and spatial segregation as key factors shaping coexistence in mixed populations.

## Introduction

Small intertidal crustaceans of the *Jaera albifrons* species complex represent perhaps the best-studied complex of sibling species among isopods. The only reliable criteria for species identification in the complex are secondary sexual characters of mature males, which consist of setae and projections on certain joints of the pereiopods (*1–3*). These traits are subject to intraspecific, interpopulation, and age-related variability (*4–8*), however, a certain characteristic pattern is constant within each species. Hybrid males, whose morphotype is known from laboratory crosses, are typically rare in nature, despite the fact that the different species often coexist (*3*, *9–11*).

While the levels of hybridization and the factors of ecological differentiation of the species of the group that prevent the formation of hybrids are largely understood (*12*, *13*), the evidence of the resulting effectiveness of the gene flow barriers between sympatric populations of the different species remains contradictory. While most species of the complex could be differentiated based on allozyme frequencies by (*14*), such a differentiation was not found for Baltic populations (*15*). Analysis of mtDNA variation the species showed no separation of the haplotypes into species-specific groups (*16*) indicative of past or ongoing introgression. Similarly, variation at microsatellite loci demonstrates that introgressive hybridization leads to gene flow between sympatric *J. albifrons* and *J. praehirsuta* in some locations, but that even in such settlements several loci maintain a strong differentiation between the morphospecies (*17*). One of the key roles in directing the introgressive hybridization and genetic differentiation between the species might belong to habitat (dis)continuity which is variable across species and geographical regions (*18*, *19*).

In the present pilot study, we aimed at analyzing the variation in the rRNA gene cluster and its differentiation between the species of the *J. albifrons* complex. Our motivation for this choice stems from the fact rRNA genes, in contrast to mtDNA and single-copy nuclear loci, are present in the genome in large numbers of copies which undergo concerted evolution which leads to gradual homogenization of the copies (*20*, *21*). Multi-copy genes under concerted evolution have larger effective population sizes (*22*) and we hypothesized that this might lead to lower fixation rates of alleles introduced by sporadic hybridization in comparison to single-copy markers.

We primarily focus on populations from the White Sea and investigate the degree and factors of spatial isolation of the species and its relationship to hybridization. We use the opportunity to also provide a first glimpse into historical relationships between the species of the complex and other members of the genus.

## Materials and methods

### Main sampling locations

Three mixed settlements were selected for spatial distribution analysis: two on Ryazhkov Island and one on Oleny Island in the Kandalaksha Bay of the White Sea (Fig. 1). Most of the material used for genotyping was collected on Ryazhkov Island (Fig. 1B), and in addition, several samples from other parts of the White Sea, as well as from the Barents, Baltic and North Seas, were used (Fig. 1A, Suppl. Table 1). Most of the material from the Oleny, Ryazhkov, and Bolshoy Solovetsky Islands was collected in June 2007, June 2008, and July 2009, respectively. Three most widespread species of the *J. albifrons* complex, out of five, were found in the studied settlements: *J. albifrons* Leach, *J. ischiosetosa* Forsman, and *J. praehirsuta* Forsman. Species of the genus *Jaera* from other morphological groups, as well as two other asellote species, were used for comparative and phylogenetic analyses (Suppl. Table 2). Most of the material was fixed in cold 96% alcohol and stored at –4 to –20°C for several days to avoid DNA degradation (*23*).

**Figure 1.**
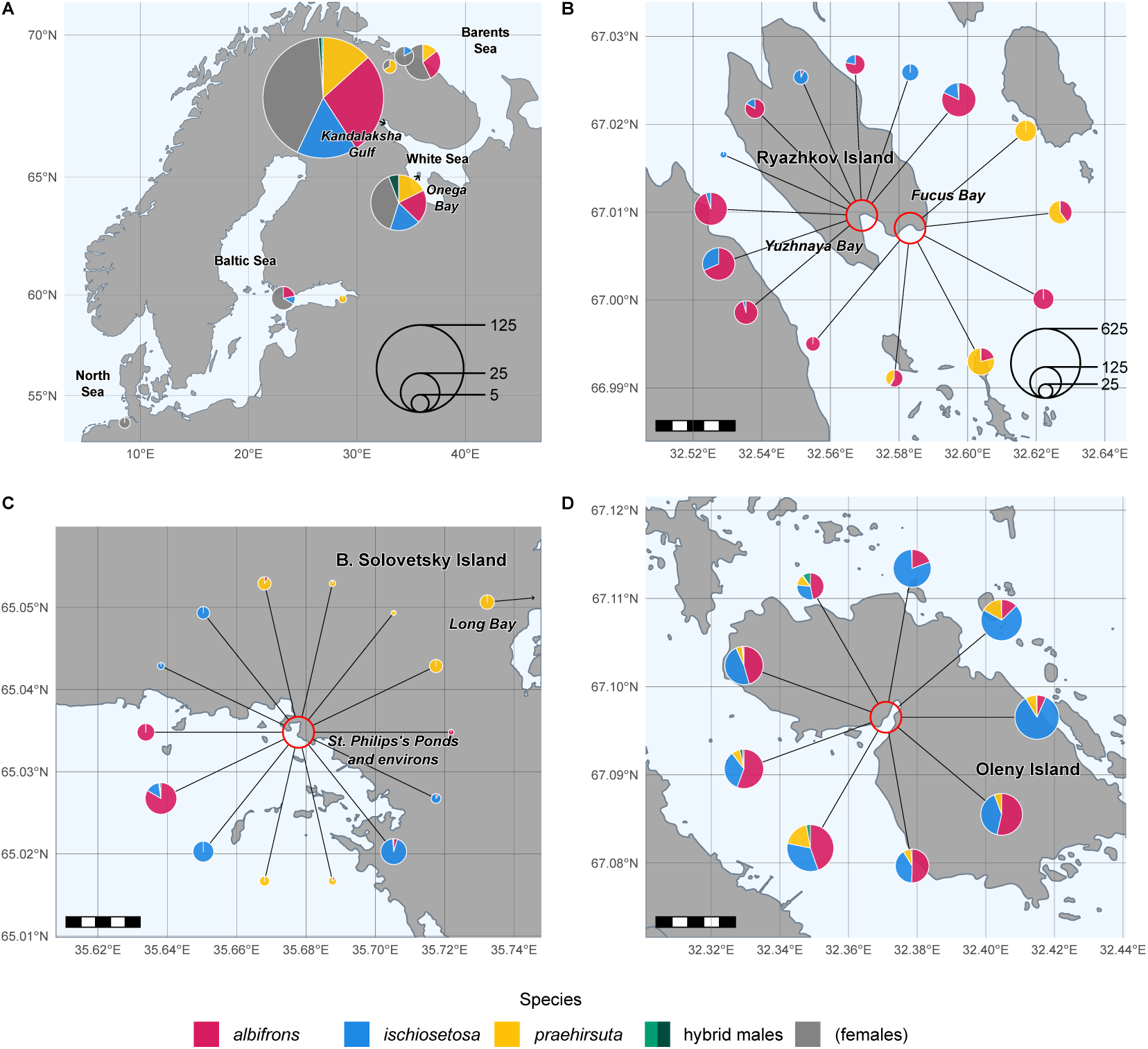
Sampling locations. (A) Map of Northern Europe showing areas from which individuals of the *J. albifrons* complex were sampled for genotyping. Pie charts indicate the proportions and number of individuals according to sex and species. (B) Location of the two settlements on Ryazhkov Island (Kandalaksha Gulf, White Sea) for which the ecological factors defining the distribution of the three *Jaera* species were studied in detail and from which most of the genotyped individuals originated. Pie charts show proportions of males of the different species in samples grouped by sampling site and substrate. (C) Location of the settlements on Bolshoy Solovetsky Island (Onega Bay, White Sea) which were used for genotyping. Pie charts show proportions of males in individual samples. (D) Location of the settlement from Oleny Island (Kandalaksha Gulf, White Sea) which was studied for potential segregation of the *Jaera* species among fucoid species. Pie charts show proportions of males in samples grouped by sampling site and fucoid species. Scale in (B-D): 1 km. Underlying map data in (A) originate from Natural Earth data (public domain) and in (B-D) from OpenStreetMap (under Open Data Commons Open Database License, https://www.openstreetmap.org/copyright).

### Collection and processing of material for genotyping and sequencing

For genotyping, crustaceans were collected from macrophytes, rocks and mussels in the intertidal zone, with several individuals per location within a radius of ∼2 m. In the Baltic Sea and Lake Mogilnoye, samples were collected by scuba diving. DNA was extracted with Chelex 100. For genotyping, each individual was dissected lengthwise before extraction and one half was dried and homogenized in 80 µl of SQ water and 12 mg of Chelex 100. Homogenized samples in Chelex were then incubated at 95°C for 20 min with periodic vortexing.

### Polymerase chain reaction

Primers for PCR were designed manually. Primer Premier v. 5.0 (Premier Biosoft Intl) and FastPCR v. 5.4.4 (*24*) were used to analyze potential dimers, annealing temperatures, and compatibility. The reaction mixes included: SibEnzyme Taq Buffer 1x, betaine 0.5 M, dNTPs 0.20–0.25 mM, SibEnzyme Taq Pol 0.05 u/μl, primers usually 0.5 μM each. 0.5–1.0 μl of DNA extraction was used per reaction. The reaction volume was 10–40 μl depending on the required amount of PCR product. The temperature profile was varied depending on the product size and primer annealing temperature. For most reactions, 35 amplification cycles were used. To obtain fragments for analyses of indels, the annealing temperature was 44°C, the elongation temperature was 65°C, and the final round of elongation was extended to 20 minutes.

### Genotyping

Two regions of the rRNA gene cluster were selected: the first transcribed spacer (ITS1) and the D1-D2 fragment (expansion segment) of the large subunit rRNA gene. An initial screen of variable positions was performed by amplifying and sequencing the rDNA regions without cloning. Primers were designed based on sequences available in GenBank and our own preliminary sequencing results. ITS1 sequences were amplified with primers JISF and 5p8aR. Complete ITS1 sequences could be obtained only for three *J. ischiosetosa* individuals. For amplification of the D1-D2 fragment, isopod-specific primers (D12FN and D12RN) were designed as the universal primers (*25*) systematically amplified protist DNA. Seven complete and four partial nucleotide sequences were obtained for the D1-D2 fragment (see Suppl. Table 1). Most of the amplicons were sequenced at the Laboratory of Molecular Systematics, Zoological Institute, Russian Academy of Sciences, on a 3130 Genetic Analyzer (Applied Biosystems) using the BigDye Terminator 3.1 kit. In the presence of intra-individual indel variability, the sequences were edited manually to discriminate the signal from two product variants.

Three indel positions: S842 and S1187 in ITS1 and D643 in D1-D2, were genotyped by amplifying the corresponding regions with one of the primers in pair labelled with Cy5 (Fig. 2A, Suppl. Table 3). The reactions for sites S842 and S1187 were performed in a single tube. The resulting PCR products were separated on ALFexpress II sequencer (Amershan Pharmacia Biotech) and the product length variation was analyzed using ALFWin Fragment Analyzer v1.03.01 (Amershan Pharmacia Biotech). Fragment sizes were calibrated using internal controls (95 and 195 bp) added to each lane (Fig. 2B). The relative amount of DNA for a peak was determined based on its height (Fig. 2C). The substitution at position D545 in D1-D2 was analyzed AluI (AG|CT) digestion of the PCR product obtained with primers D12RN and JD426. Digestion products were visualized by vertical electrophoresis in a 5% polyacrylamide gel and silver-staining. The resulting images were analyzed using TotalLab v 2.01 (TotalLab Ltd) (see Fig. 2A). If two alternative restriction products were present simultaneously, the total relative DNA amount for the corresponding bands was determined based on their staining intensity. The total staining intensity was found to be a more accurate measure of restriction product ratios than peak height for this genotyping approach.

**Figure 2.**
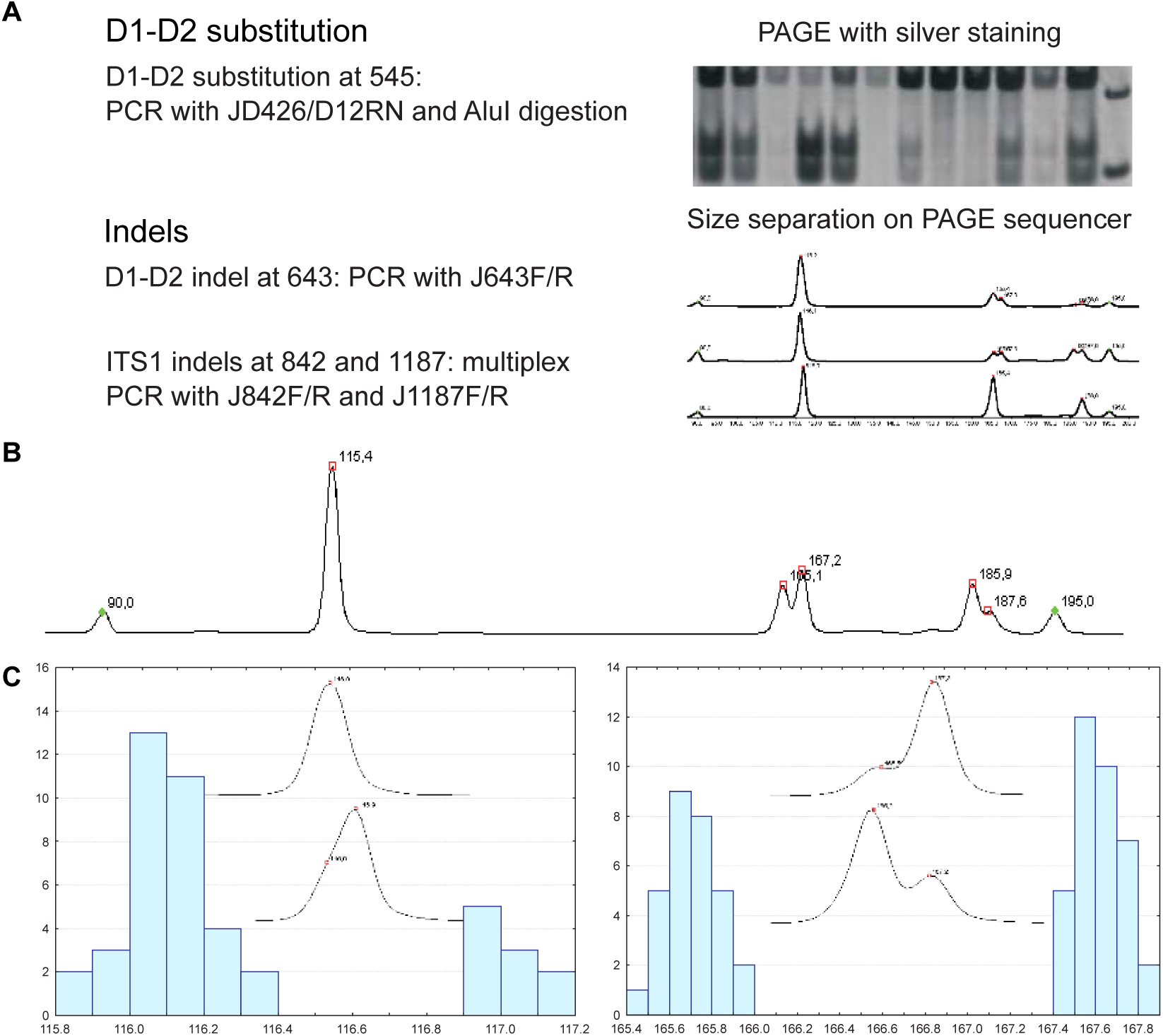
Genotyping strategy for variable positions in the ITS1 and D1-D2 fragments in *J. albifrons* complex. (A) The four variable positions were analyzed by amplifying the corresponding regions and analyzing the lengths of the fluorescently labelled amplicons on gel sequencer (indels) or analyzing digestion products on gel electrophoresis with silver staining (substitution). (B) Example electropherogram for a single individual showing intraindividual variation at the indel sites. (C) Example peaks and frequency distributions of the estimated fragment sizes for ITS1 sites S1187 (left) and S842 (right) for a subset of the analyzed samples.

Each individual was thus characterized by the relative amount of the corresponding alternative variants. For many individuals, genotyping was performed multiple times to assess reproducibility by either repeating PCR reactions or only electrophoresis. Genotyping at site S1187 was the least reliable due to the difficulty of determining the presence and height of the minor peak (see Fig. 2C). For this reason, electropherograms for this site were verified several times.

### Collection and processing of material for spatial distribution analysis

Quantitative sampling for spatial distribution was performed using three different schemes at each site.

For the settlement in Fucus Bay on Ryazhkov Island, an analysis of the possible vertical distribution of the two local *Jaera* species was conducted. A series of preliminary samples was used to determine that two *Jaera* species coexist in the upper part of the fucoid belt. Next, two transects separated by 15 m were selected, along which samples were taken at three levels. At each level within a transect, three thalli of *Fucus vesiculosus* were collected (Fig. 3A).

**Figure 3.**
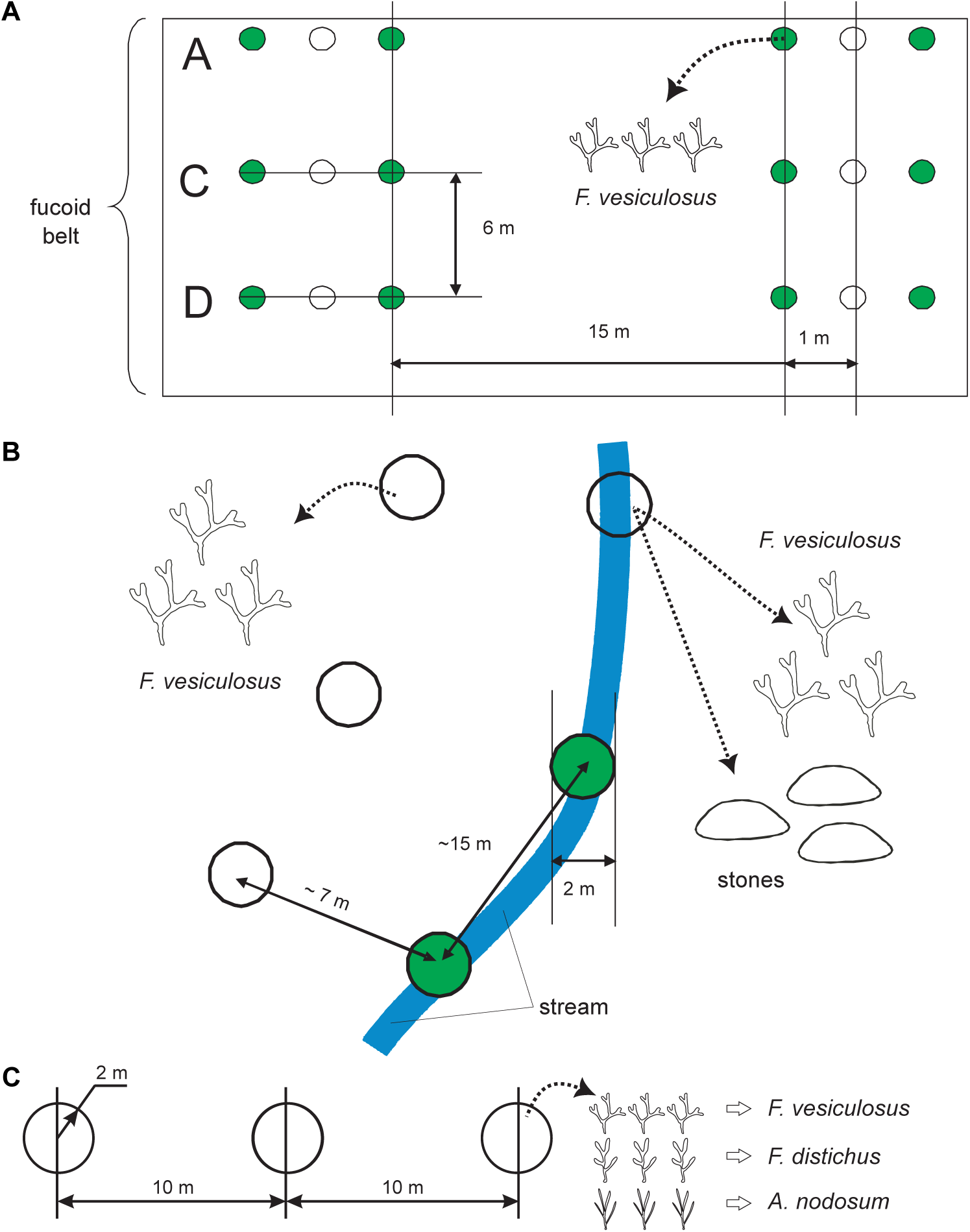
Sampling design for the analysis of spatial distribution of the three species of the *J. albifrons* complex in the two settlements on Ryazhkov Island: Fucus Bay (A) and Yuzhnaya Bay (B), and on Oleny Island (C).

For the mixed settlement in the Yuzhnaya Bay on Ryazhkov Island, the influence of two interrelated factors was studied. Here, a stream flows into the corner of the bay and at low tide flows through the intertidal zone. Therefore, the sampling design for this settlement included an analysis of the influence of the substrate on the distribution of the *Jaera* species in the stream, as well as the influence of the stream itself on the distribution of *Jaera* on fucoids. In the intertidal part of the stream, three *Fucus vesiculosus* thalli and three groups of stones were collected at three sites along the stream flow. Using a similar scheme, samples were taken just outside the stream from fucoids (Fig. 3B).

For the settlement from the Oleny Island in which all three *Jaera* species were known to occur, we tested the hypothesis of possible segregation of the different species on different fucoid species. For this purpose, three sites were selected along a transect in the fucoid belt (Fig. 3C). At each site, three thalli of three fucoid species (*Fucus vesiculosus*, *Fucus distichus*, and *Ascophyllum nodosum*) were randomly selected within a 2-meter radius.

In the laboratory, all isopods were washed off the fucoids and stones. The thalli were weighed after removing the surface water. Fucoid surface area for the samples from the settlement on the Oleny Island were identified based on a series of preliminary measurements of a sample of weighed thalli: *Fucus vesiculosus*: *S* = 9.5*M* + 165.3 (*R^2^* = 0.43), *F. distichus*: *S* = 15.3*M* + 34.4 (*R^2^* = 0.89), and *Ascophyllum nodosum S* = 13.8*M* + 33.3 (*R^2^*= 0.58), where *S* is the surface area in cm^2^ and *M* is mass in grams.

Males were identified to the species level based on secondary sexual characteristics. For each sample, the ratio of males of the different species was determined (see Fig. 1B,D for summary), as well as their population densities which were used as dependent variables in the analysis of variance. For the samples from the Oleny Island, the density of males of each species was determined relative to the calculated thalli surface area, and for the Ryazhkov Island settlements, relative to the mass of *F. vesiculosus* thalli. To analyze the spatial distribution of females based on genotyping results, samples from settlements on the Ryazhkov Island were used. The same samples were used, but the experimental design was modified. Seven genotyped females were collected from two transects at each of the three levels in Fucus Bay. These females were obtained from the two most distant fucoids within a horizon (see Fig. 3A). In the Yuzhnaya Bay, seven genotyped females were collected from each substrate type (rocks and fucoids) at two of three sites in the stream (see Fig. 3B). In these cases, quantitative genotyping features for each female were used as features for the (multivariate) analysis of variance (ANOVA) (see below). Similar random samples of males were drawn as controls.

### Statistical analysis

To identify factors influencing the spatial distribution of the *Jaera* species, we used analysis of variance with GLM in SPSS Statistics v. 19.0.0 (SPSS Inc.) and multivariate analysis of variance in PERMANOVA (*26*, *27*). Post-hoc tests included Tukey test and multiple *t*-test, respectively. The applicability of the analysis of variance for univariate analysis was verified in Statistica v. 7.0 (StatSoft Inc.) using the Cochran test, and for the multivariate analysis, in PERMDISP using the multivariate analogue of Levene’s test (*28*). In cases of significant heteroscedasticity, *arcsin*- and ∜-transformations were used for the corresponding variables. Multidimensional scaling (MDS) was used to represent genotyping data, and ANOSIM was used to compare multivariate distributions. Both analyses were performed in Primer v. 5.2 (Primer-E Ltd).

To determine the species identity of females from the Ryazhkov and B. Solovetsky Island settlements, discriminant analysis in Statistica was used to predict species based on rRNA genotypes. Males from these settlements served as the training set, and the effectiveness of the discriminant functions was tested based on their ability to identify males a posteriori.

Differences at a significance level of *p* ≤ 0.05 were considered significant. ggplot2 v. 4.0.1 (*29*), pheatmap v. 1.0.12 (*30*), as well as standard Statistica tools were used to plot the graphs. Spatial data were visualized with the help of Simple Features v. 1.0-23 (*31*) with the maps originating from Natural Earth data (https://www.naturalearthdata.com/, public domain) and OpenStreetMap (https://www.openstreetmap.org/, under Open Data Commons Open Database License).

### Phylogenetic analysis

Phylogenetic analysis of the relationships among *Jaera* species was performed based on fragments of the rRNA gene cluster: 3’ end of the SSU gene, the D1-D2 fragment of the LSU gene and ITS1 (Suppl. Table 4). The SSU fragment was amplified with primers 1800 and 18a1 and sequenced with primer 1800 and the D1-D2 fragment was amplified with primers D12FN and D12RN and sequenced in both directions (see Suppl. Table 3). *Asellus aquaticus* was used as an outgroup for rooting the trees, and representatives of other Janiroidea were taken as closer outgroups: *Carpias*, *Ianiropsis*, and two species of the Munnopsididae (see Table 8).

Alignments were performed using Muscle (*32*) with default settings. The ITS1 alignment proved to be too variable and was not used for phylogenetic analysis (see below). The final alignments for the three datasets were 525 bp for the D1-D2 fragment, 592 for the 3’ fragment of the SSU gene. Maximum likelihood and maximum parsimony trees were searched for with PAUP* v. 4.0b10 (*33*) with bootstrap analysis (1000 permutations). MrBayes v. 3.1.2 (*34*) with default settings was used for the search of the optimal Bayesian tree. Suitable nucleotide substitution models were determined in Modeltest according to the Akaike information criterion (AIC) (*35*).

*ITS1 analyses*. For secondary structure analysis of ITS1, individual nucleotide sequences were analyzed using the Mfold v. 3.2 webserver (*36*). The results were visualized in Varna v. 3.7 (*37*). Sequence simplicity and repetitiveness were analyzed with Simple v. 4.0 (Medical Research Council Harwell) and Phobos v. 3.3.12 (imperfect search strategy, repeat length 1-100, repeat identity 90-100%, typical score constraint) (*38*). dotplot from EMBOSS v. 6.6.0.0 (*39*) was used to analyze large duplications in ITS1 sequences.

## Results

Two locations in the White Sea (Fig. 1) were chosen for analysis of factors defining spatial distribution of the different *Jaera* species: Fucus Bay and Yuzhnaya Bay of Ryazhkov Island and a mixed settlement on Oleny Island. The settlements from Fucus Bay and Yuzhnaya Bay were the primary focus of the rRNA genotyping effort, with additional material recruited from B. Solovetsky Island, locations from the Barents Sea and limited material from the Baltic and North Sea (see Fig. 1).

### Variability of the ITS1 and D1-D2 regions

Two regions of the rRNA gene cluster, ITS1 and D1-D2, were chosen as genes known to demonstrate high variability in other cases of closely related species. Complete or partial D1-D2 sequences were obtained for a total of 11 individuals of *J. albifrons*, *J. ischiosetosa* and *J. praehirsuta* from the White and Barents Seas. Importantly, no cloning was performed which often resulted in mixed sequences in Sanger sequencing. Only two sites were found to be variable across species and thus chosen for genotyping (Table 1). A third position was variable among the sequenced *J. praehirsuta* individuals (see Table 1).

**Table 1.**
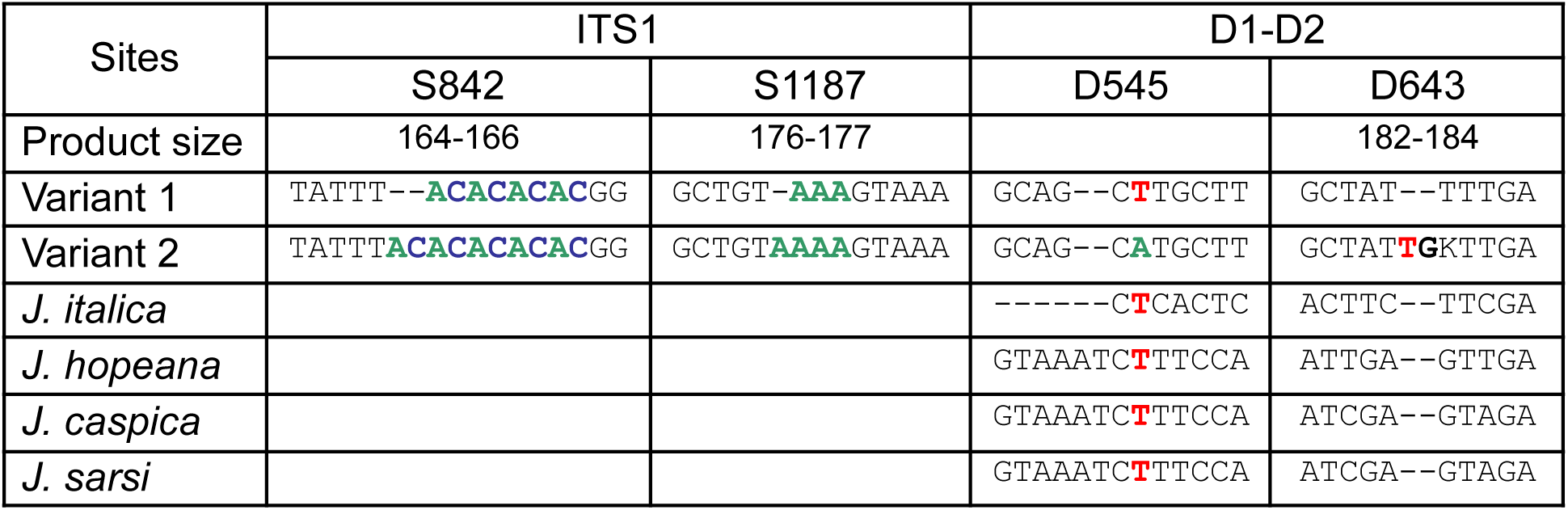
Variable positions in the rRNA gene cluster used for genotyping. Homologous regions from other *Jaera* species are provided for comparison (in *J. cornuta*, these regions are partially deleted). The K ambiguity next to indel 643 reflects additional variability in individuals without the insertion.

Overall, the variability of the D1-D2 fragment in the *Jaera* genus was found to be very high as it exceeded the variability among the Munnopsididae and was comparable to the intergeneric variability between *Carpias* and *Ianiropsis* (see *Phylogeny* below).

ITS1 was more variable in the *J. albifrons* group than the D1-D2 segment. Sequencing was attempted for eight individuals, yet due to widespread intra-individual variability in short indels, the spacer could be fully sequenced only for three *J. ischiosetosa* males (see Suppl. Table 1). This does not allow us to draw conclusions about the general ITS1 variability for the *J. albifrons* group species, however interspecific variation was detected for at least six sites based on the partial sequences (Suppl. Table 5). Five of these sites are substitutions and short indels, two of which were chosen for genotyping. The sixth variable position is a region of 22–23 bp with multiple indels and substitutions (see Suppl. Table 5). This site demonstrated only two alternative variants and did not show a particular species or geographical association, and intra-individual polymorphism was registered in one individual.

Interestingly, the complete ITS1 sequences from *J. ischiosetosa* showed the presence of a large duplication (Fig. 4A) with two homologous regions 288 bp (C1) and 279 bp (C2) separated by a 35-nucleotide region (Fig. 4B). These two regions are 91.0% identical and are predicted to have a specific secondary structure. The ITS1 indels analyzed on the bulk material (see below) belong to two homologous hairpins within the duplicated fragment (Fig. 4C).

**Figure 4.**
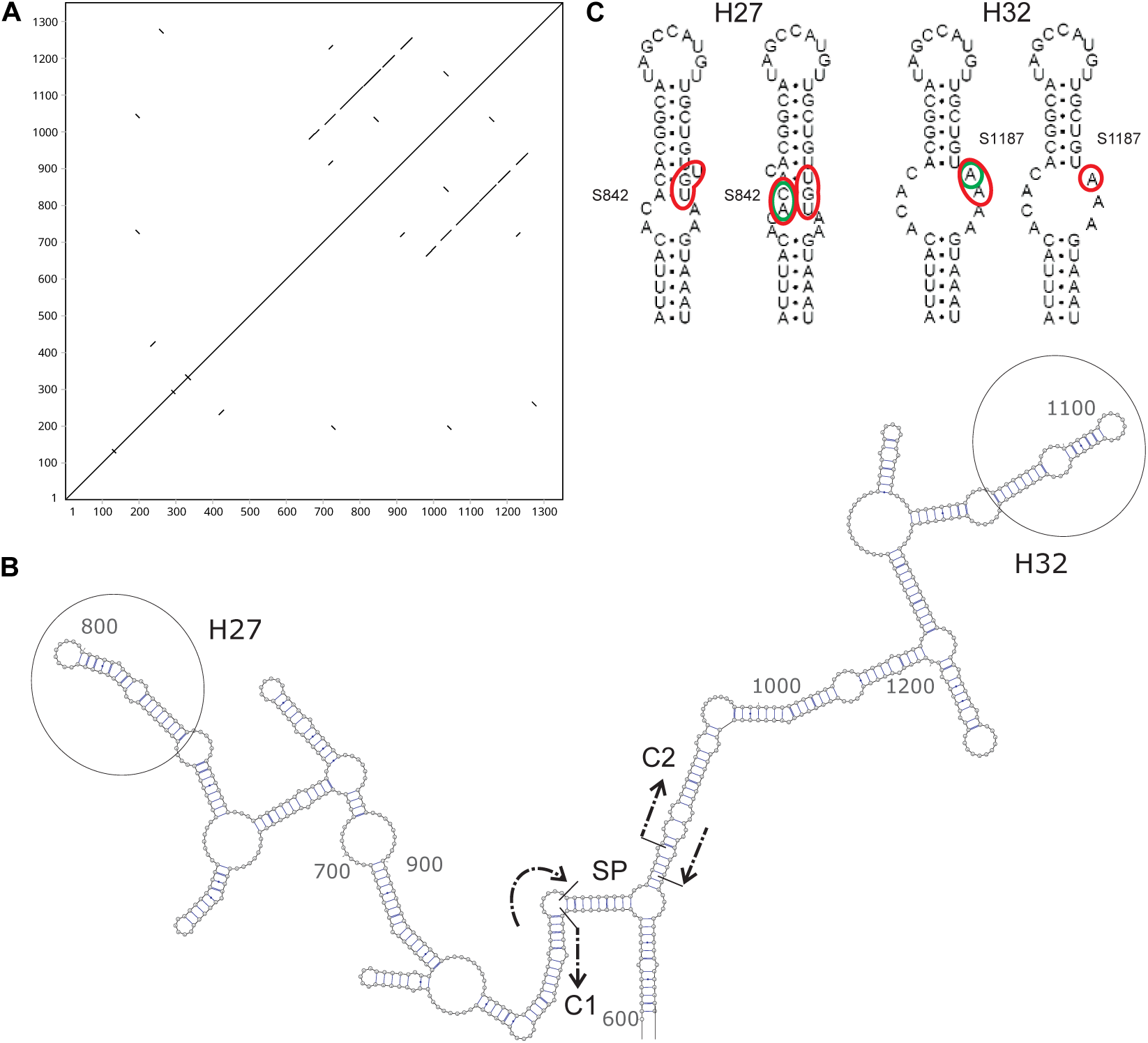
Structure of the ITS1 sequence in the *J. albifrons* complex. (A) Self-similarity dotplot (window size of 10 bp). (B) Fragment of the most stable secondary structure of *J. ischiosetosa* ITS1 (*E* = –281.99 KCal/Mol) around the duplicated region. Arrows mark the boundaries of the two homologous regions (C1 and C2) and the 35-nucleotide spacer between them (SP). H27 and H32 are two homologous hairpins with variable positions. (C) Closeup of the hairpins H27 and H32, each with two alternative variants observed in *J. albifrons* species complex. Positions differing between the two hairpins are highlighted in red and positions variable between different alleles (labelled accordingly) highlighted in green.

At a higher taxonomic level, ITS1 was also more variable than D1-D2. No blocks of confident homology across Asellota could be identified. In addition, ITS1 of different species varied significantly in length and GC content (Table 2). Analysis of linguistic complexity of ITS1 in Asellota showed a relatively low but significant linguistic simplicity of these sequences, although generally only a small fraction of the spacer was occupied by tandem repeats (see Table 11).

**Table 2.**
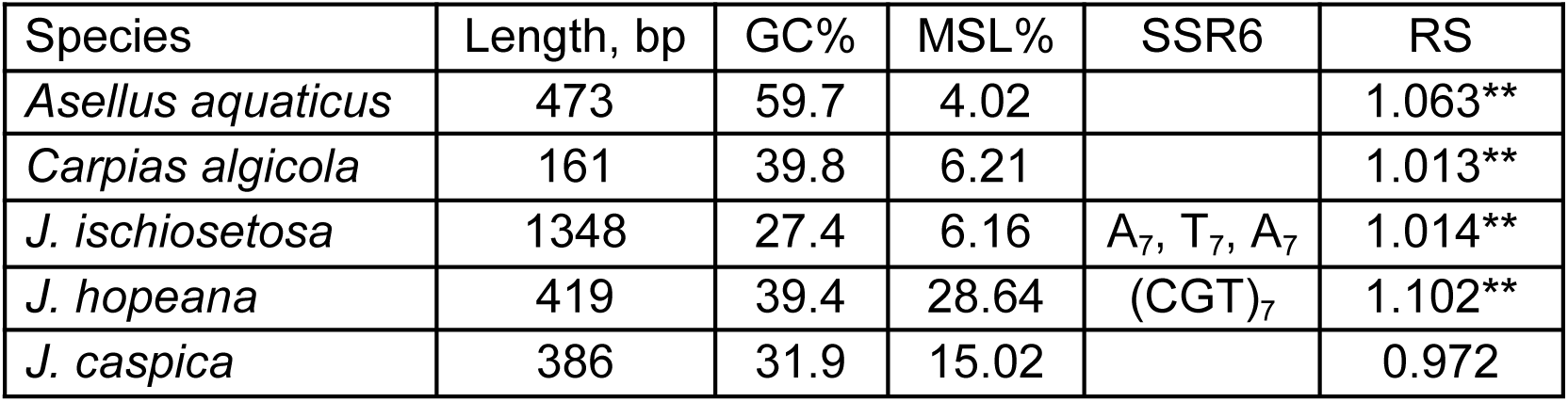
Main characteristics of ITS1 sequences from Asellota. RS – relative simplicity of the sequence. Values significantly different from RS for random sequences (*p* ≤ 0.01) are indicated with **. MSL% – fraction of microsatellites as a percentage of the total ITS1 length; SSR6 – short sequence repeats with a number of units >6. The boundary between ITS1 and 5.8S in *J. ischiosetosa* is not defined and the sequence likely also includes ∼10 bp of the 5’ end of 5.8S.

### Phylogeny

To analyze the phylogenetic relationships between species of the genus, representatives of all morphological and biological groups were collected except the unusual deep-sea *J. tyleri* Brandt et Malyutina, 2014 in Linse et al 2014 (*40*). For reconstructions, we used two regions of the rRNA cluster: a 3’ fragment of the SSU gene and the D1-D2 segment of the LSU gene. Given the hypervariability of the ITS1 sequences (see above), it could not be used as a phylogenetic marker. The obtained trees were concordant between D1-D2 and SSU but differed sharply from the morphology-based reconstruction by (*41*). Thus, the Mediterranean group of species (represented here by *J. cornuta* Karaman [= *J. normanni* forma *cornuta* Karaman, *J. massiliensis* Lemercier] and *J. italica* Kesselyák) appeared paraphyletic with respect to the *J. albifrons* complex (the Atlantic group) with *J. italica* being closest among the sampled species to the *J. albifrons* complex. (The sequence AF279610 (*23*) appears to belong to an individual from the *albifrons* complex and not *J. nordmanni* as it is almost identical to the other sequences from the complex and does not show similarity with the SSU sequence from *J. cornuta* representing the *J. nordmanni* complex in our sample). Analogously, *J. hopeana* demonstrated a sister-group relationship with the Ponto-Caspian group (*J. sarsi* and *J. caspica*) thus contradicting its basal position within the genus. This indicates that its placement in a separate subgenus *Jaera* (*Metajaera*) Verhoeff, 1943 is unjustified.

**Figure 5.**
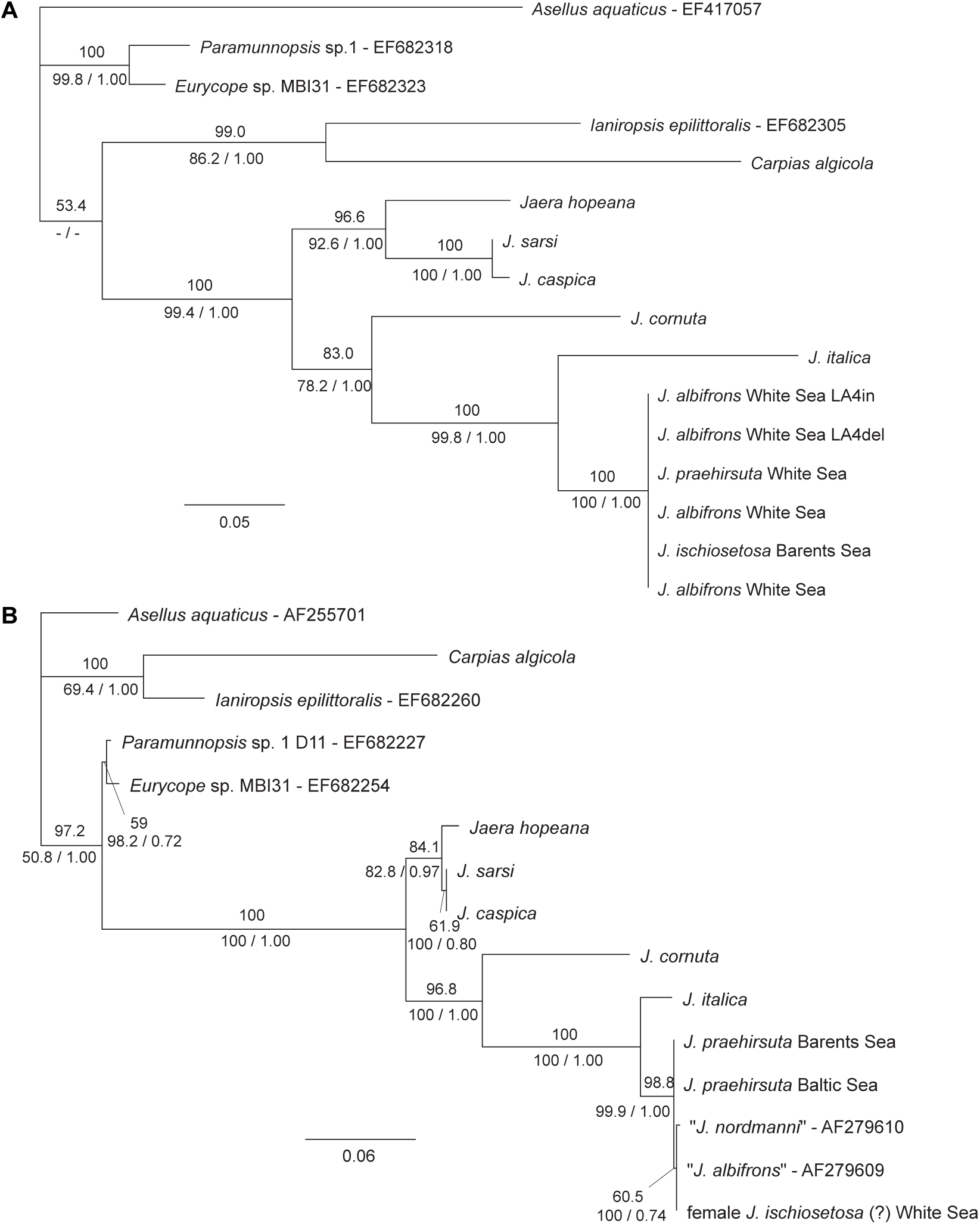
Phylogenetic position of the *J. albifrons* complex. (A) Maximum likelihood (ML) phylogenetic tree based on the 525-nt nucleotide alignment of the D1-D2 fragment of LSU under the GTR+G+I model (*α* = 1.961, *p_inv_* = 0.247, four categories). (B) ML tree based on the 592-nt nucleotide alignment of a fragment of SSU under the GTR+G (*α* = 0.294; four categories). Numbers above the branches indicate ML bootstrap support values (*n* = 500), numbers below the branches indicate maximum-parsimony bootstrap support values (*n* = 1000) and posterior probabilities from the Bayesian inference, respectively. Both trees are rooted by the sequences from *Asellus aquaticus*.

### Genotyping

Variation at four sites within the rRNA gene cluster was analyzed for samples of *J. albifrons* complex from the White Sea and other locations by amplifying the corresponding regions and assessing relative amounts of alternative variants (see Fig. 2). Only 64 out of 310 genotyped individuals showed a single variant at each of the four sites. In the remaining individuals, one or more sites demonstrated both alternative states (Fig. 6). Clustering the individuals based on the resulting quantitative genotypes, revealed the presence of: 1) a cluster dominated by *J. ischiosetosa* which demonstrated the lowest levels of intraindividual variation and was most dissimilar to the other individuals, 2) a cluster dominated by *J. albifrons*, 3) a cluster with most of the *J. praehirsuta* individuals but with a sizeable proportion of *J. albifrons* males and 4) a separate cluster which encompassed the few genotyped individuals from the Baltic Sea, including representative males of all three species, as well as several individuals from the White and Barents Seas. Females did not form a separate cluster in this analysis thus indicating that the rRNA variants are not sex-linked. A similar pattern was replicated by MDS (Fig. 7A). The cloud represented by males of *J. ischiosetosa* and associated females was most distinct with many individuals sharing the same genotype. *J. praehirsuta* formed a separate cloud with least similarity to *J. ischiosetosa* but overlapping the cloud of *J. albifrons*. This last species was found to be most diverse with hints of geographical variation as indicated by the differences between the populations from the White Sea and the Barents Sea (Fig. 7). The few analyzed individuals from the Baltic Sea fell in a separate cloud close to *J. albifrons* individuals from the Barents Sea.

**Figure 6.**
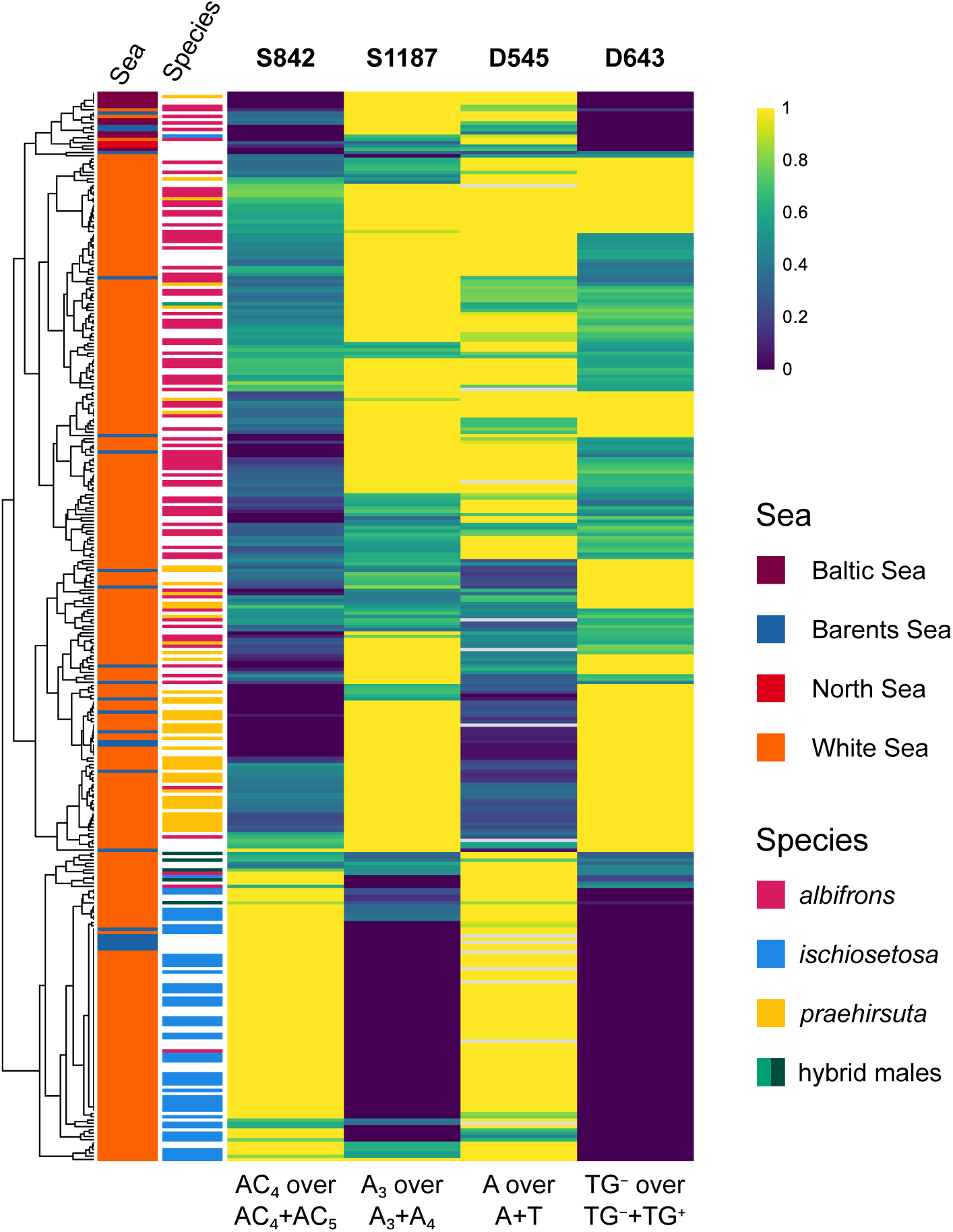
Summary of genotyping results for four positions found to be variable among members of the *J. albifrons* complex. The heatmap is based on the proportion of one of the two alternative variants at the corresponding position. S852 and S1187 are variable positions in the ITS1 sequence and D545 and D643 fall in the D1-D2 region. Complete-linkage was used to cluster the individuals. Individuals without species identification are females.

**Figure 7.**
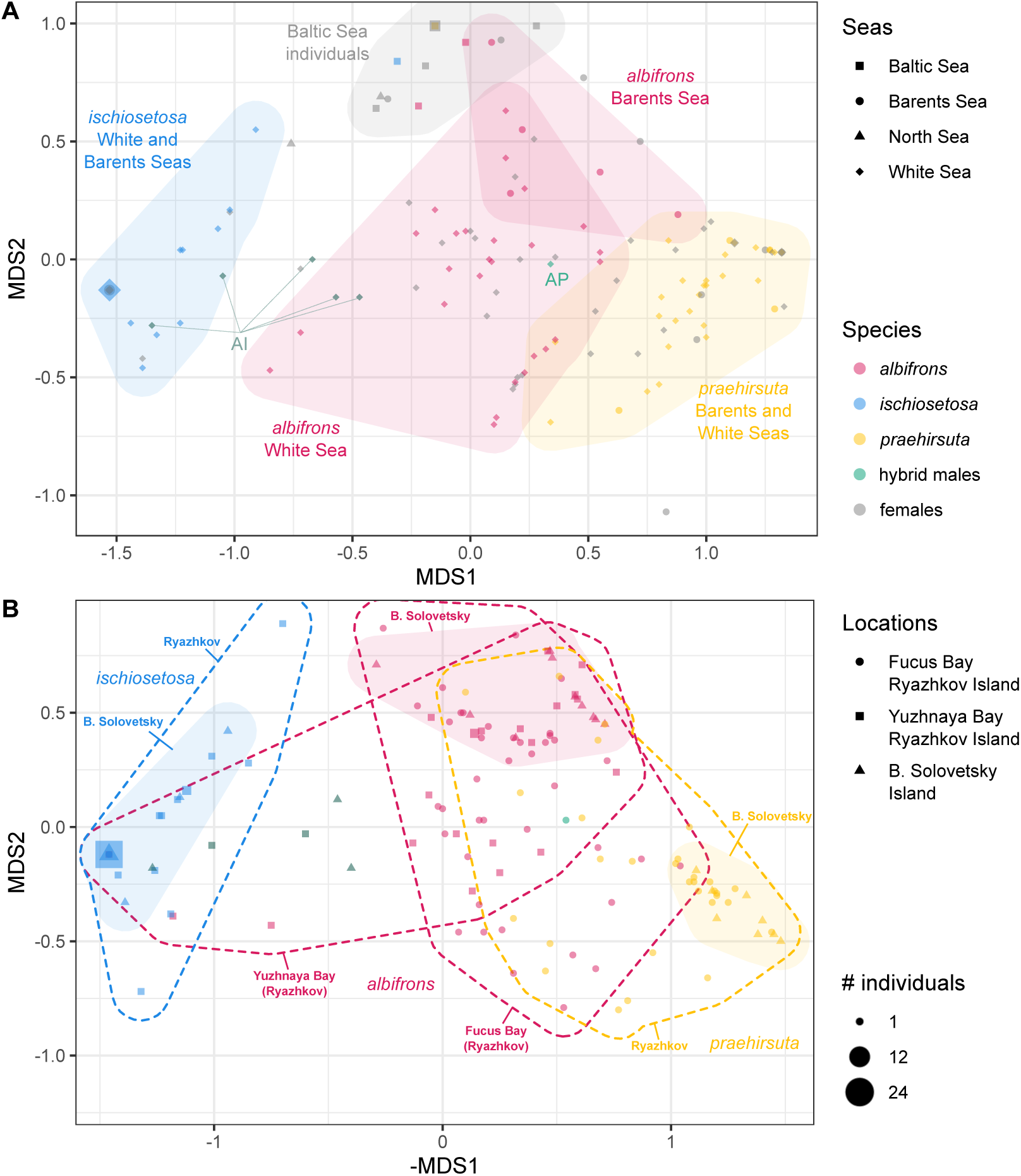
Ordination of individuals of different *Jaera* species along the MDS axes based on the variation at four sites in D1-D2 and ITS1. (A) MDS analysis for individuals originating from diverse locations with Ryazhkov Island populations represented by random samples of 15 males of each species and 30 females. Individuals with identical genotypes are combined and the size of the points reflects the number of the agglomerated individuals. (B) MDS analysis for individuals from the White Sea. Only males are shown. Filled areas unite individuals from B. Solovetsky Island and dashed lines unite individuals from the two settlements on Ryazhkov Island. Females are not shown for clarity. Color scheme and point sizes apply to both panels.

Males identified by morphological criteria as hybrids between *J. albifrons* and *J. ischiosetosa* were located between the clouds of these two species, and two of the five analyzed males appeared indistinguishable from *J. ischiosetosa* males. The only analyzed *albifrons*-*praehirsuta* hybrid grouped among *J. albifrons* individuals.

The general trend seen on the larger geographic scope was found to be to a large degree valid for material from two closely located mixed settlements of Ryazhkov Island which were analyzed in more detail (Fig. 7B). Here, males of *J. ischiosetosa* and *J. praehirsuta* formed two independent clouds, and males of *J. albifrons* located between them and significantly intergrading with the cloud of *J. praehirsuta* males. ANOSIM could not distinguish between the two subpopulations of *J. albifrons* from the Fucus Bay (where it is sympatric with *J. praehirsuta*) and from the Yuzhnaya Bay (sympatric with *J. ischiosetosa*) (*R* = 0.044; *p* = 0.17), although there was a trend for the *J. albifrons* males from the Fucus Bay to be more similar to *J. praehirsuta* and similarly, among the *J. albifrons* males from the Yuzhnaya Bay we found two males falling in the cloud of *J. ischiosetosa*. Despite the broad overlap, ANOSIM for the Fucus Bay did reliably distinguish between males of the *J. albifrons* and *J. praehirsuta* (*R* = 0.431; *p* = 0.001). The much smaller sample of males from B. Solovetsky Island showed a complete separation of the clouds corresponding to three species (see Fig. 7B).

### Appearance of hybrid males

Hybrid males carrying secondary sexual characteristics of more than one species were found in all four locations from the White Sea (Table 3). Among the three possible combinations of hybrids, only hybrids for which one of the parent species was *J. albifrons* (*albifrons*-*ischiosetosa* and *albifrons*-*praehirsuta*) were found, even on Oleny Island, where all three species were found in the same samples. Overall, hybrid frequencies did not exceed 1% for most settlements. However, in the settlement on Oleny Island, the frequency of the *albifrons*-*praehirsuta* hybrids was exceptionally high (∼3%), with average of per-sample proportion of hybrid males of 7.72% ± 6.279 (*n* = 9) relative to the two parental species in samples which contained the hybrids (Suppl. Fig. 1).

**Table 3.** Hybrid males in the White Sea settlements. “AI” and “AP”: *albifrons*-*ischiosetosa* and *albifrons*-*praehirsuta* hybrids, respectively. “Proportions” indicate the proportions of hybrid males among all males, and those in parentheses include only the parental species (for three-species settlements).

| Settlement | Hybrid males/<br>total | Proportion of hybrid males (%) |  |
| --- | --- | --- | --- |
|  |  | AI | AP |
| Fucus Bay, Ryazhkov Island | 2/347 |  | 0.58 |
| Yuzhnaya Bay, Ryazhkov Island | 2/666 | 0.30 |  |
| Bolshoy Solovetsky Island | 3/444 | 0.68 (0.86) |  |
| Oleny Island | 32/1802 | 0.44 (0.50) | 1.33 (2.85) |

### Spatial distribution of the species

The quantitative ratios of the three species varied significantly between the settlements (see Fig. 1B,D). Microdistribution of the three species was investigated in detail for Ryazhkov and Oleny Islands.

On Ryazhkov Island, the two studied locations were separated by less than one kilometer (see Fig. 1B). Yet, in the Yuzhnaya Bay only *J. albifrons* and *J. ischiosetosa* were found (*n* = 666) while in the Fucus Bay — mainly *J. albifrons* and *J. praehirsuta* (*n* = 347) were identified and only a single *J. ischiosetosa* male.

For Fucus Bay, pilot sampling revealed that the level above the fucoid belt was inhabited by males of *J. albifrons*, while males of *J. praehirsuta* were found in the lower part of the fucoid belt. The quantitative experiment was thus designed to analyze the segregation of these two species in the part of the fucoid belt where they were expected to coexist (see Fig. 3A). Suppl. Table 6 shows the results of the analysis of variance for the factors “littoral level” and “site” with response variables represented by the population densities of the males of each species or the relative proportion of the species: A reliable effect for the given experimental design was found only for the factor “level” for the species proportion. Indeed, all three selected intertidal levels differed significantly from each other in the proportions of males of the two species (Fig. 8, Suppl. Table 7). In particular, in the upper part of the fucoid belt, males of *J. praehirsuta* were absent entirely.

**Figure 8.**
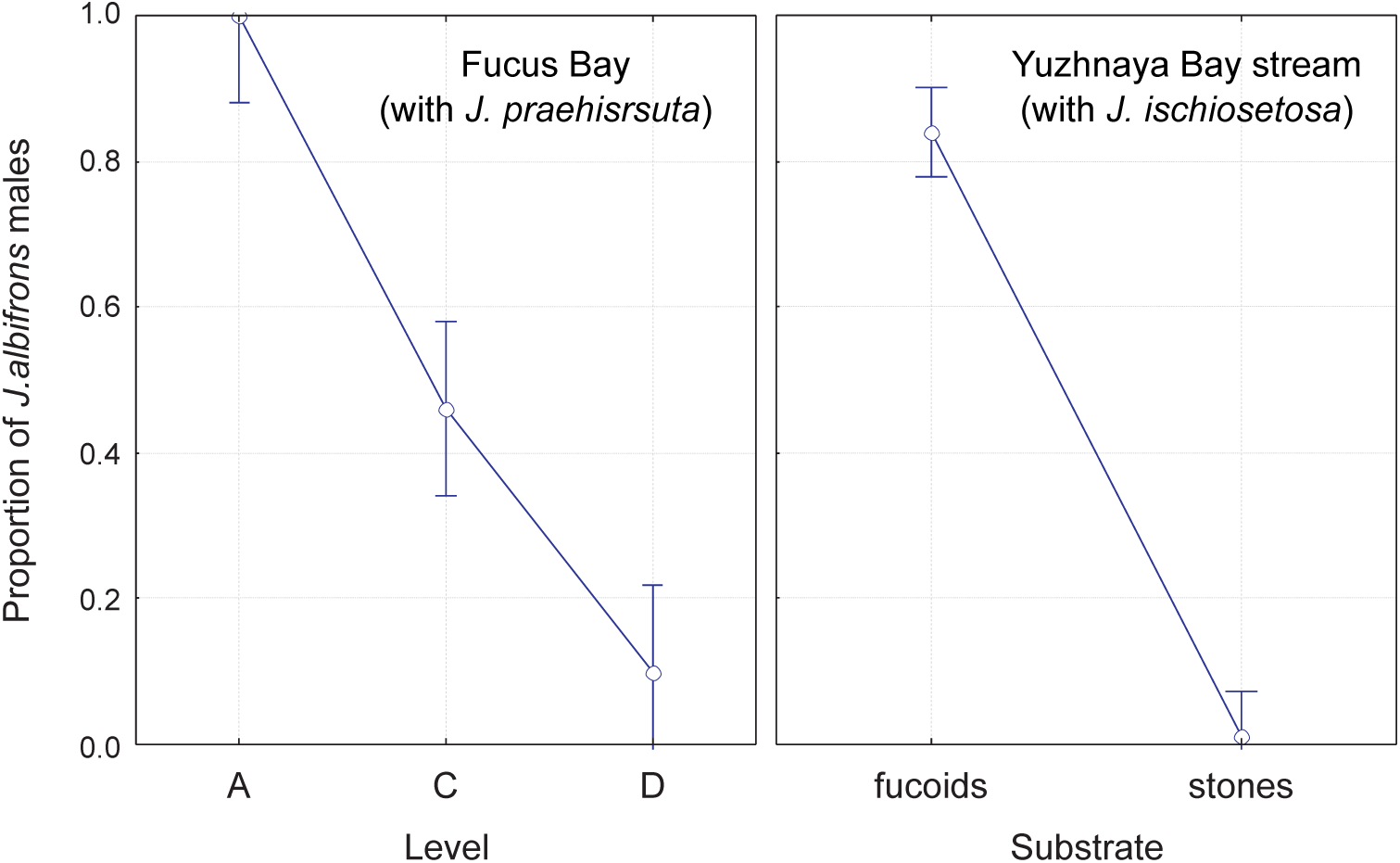
Proportion of *J. albifrons* males in the Ryazhkov Island settlements as a function of intertidal level in the Fucus Bay settlement (left) and substrate in the samples from the stream in the Yuzhnaya Bay (right). Plotted are average proportions with 95% confidence intervals.

For Yuzhnaya Bay, preliminary data showed that, on the one hand, males of *J. albifrons* were predominantly found on fucoids at a distance from the stream, while in the upper part of the intertidal zone, at the mouth of the stream in the absence of fucoids, *J. ischiosetosa* were found. Therefore, a series of samples were taken in the stream from two types of substrates and near the stream from fucoids (see Fig. 3B). Analysis of variance for the factors “stream” (distance from the stream) and “level” (distance from water’s edge at low tide) did not reveal any influence of these factors on either the population density of each of the two species or their proportions (Suppl. Table 8). At the same time, within the stream, the substrate had a significant effect on the species proportion (Suppl. Table 9), with males of *J. albifrons* absent on rocky substrates (see Fig. 8). In this case, the influence of the factor “level” (distance from water’s edge at low tide) was also insignificant.

In the settlement from Oleny Island, all three *Jaera* species were found on fucoids represented by three dominant species: *Fucus vesiculosus*, *F. distichus* and *Ascophyllum nodosum*. The sampling scheme was designed to identify possible segregation of the different species across different fucoid species (see Fig. 3C). Analysis of variance for the densities of individual species did not reveal any significant effect of the factors “fucoid species” and “site” (Suppl. Table 10). A similar analysis for the proportions of each species by these factors revealed a significant effect only for the “site” factor for *J. albifrons* and *J. ischiosetosa* (see Suppl. Table 10). Indeed, the first site differed sharply in the proportions of the males of these two species from the other two sites (Suppl. Table 11). Multivariate analysis for the frequencies of the three species also revealed a significant effect of “site” and no significant effect of the fucoid species (Suppl. Table 12).

### Spatial distribution of females

Based on genotyping characteristics and their relative species specificity, the spatial distribution of females was analyzed for two settlements on Ryazhkov Island. Multivariate analysis of variance was used to test hypotheses similar to those regarding the distribution of males based on morphological characteristics (see above) but using discriminant analysis to tell the species apart. The success of species identification was dependent on species and location with most confident identification for *J. ischiosetosa* and B. Solovetsky Island (Suppl. Table 13). For the Fucus Bay, with the given experimental design, no significant effect of “site” and “level” was detected for females, nor could it be detected for a similar sample of males (Suppl. Table 14). However, for females, the interaction of these factors was significant: in the second site, females from the two extreme levels differed significantly from each other (Suppl. Table 14). For the Yuzhnaya Bay, the influence of “site” and “substrate” was tested for samples from the stream. A significant influence of the substrate factor was found for both females and a similar sample of males (see Suppl. Table 14).

## Discussion and conclusions

In this pilot study we analyzed variation in the rRNA genes in the *J. albifrons* complex to explore whether they can be used as diagnostic tools for species identification and as phylogenetic markers. The very shallow variability of the analyzed genes in the *J. albifrons* complex stands in stark contrast to other species of the genus clearly demonstrating that these species are nascent. While the level of variability in D1-D2 was found to be negligible (three variable sites per 960 bp), ITS1 was somewhat more variable within the group (*ca*. 10 variable sites per *ca*. 1350 bp). Such variability is generally consistent with or even lower than intraspecific variability for other animal groups (*25*, *42*, *43*). In organisms studied previously, both markers (as well as ITS2) were shown to be highly variable at the interspecific level, even in very similar species (*44–47*). Exceptions are also known (*48*), when the level of variability in genetic markers (ITS2 and mtDNA in the cited work) is very shallow for a group of closely related species: in the absence of a correlation between genetic and morphological traits this is strong evidence in favor of a single freely interbreeding species. Nevertheless, this does not appear to be the case with species of the *J. albifrons* complex.

Despite the low inter-specific variability, *J. ischiosetosa* both shows the lowest intra-individual variation at the studied rRNA positions among the three analyzed species and comes close to having species-specific diagnostic variants. The most clear differentiation was observed for *J. ischiosetosa* and *J. praehirsuta* which simultaneously represent the species pair with least ecological overlap and the one for which we found no hybrid males identifiable using morphological criteria. *J. albifrons*, on the other hand, showed an overlap with *J. praehirsuta* despite differing from it statistically. Given the fact that *J. albifrons* and *J. praehirsuta* are known to hybridize introgressively (*10*, *17*), this is best explained by rRNA copies exchanged between the species and not by ancestral variation. *J. albifrons* males from the two settlements from Ryazhkov Island, while largely representing a single pool of genotypes, show signs of local hybridization with *J. ischiosetosa* and *J. praehirsuta*, respectively. Although for comparison we recruited only several individuals from Bolshoy Solovetsky Island, they appear to form well separated clouds corresponding to the three species, thus hinting at the possibility that the rate of introgressive hybridization varies between settlements in the White Sea. The material from the Baltic Sea analyzed here is very limited, but the clustering of all Baltic individuals in a single group, irrespective of morphological species, seems to support the earlier observations of lack of differences between the three *Jaera* species in Baltic populations in allozyme frequencies (*15*).

Comparison to other *Jaera* revealed that the shallow genetic variability in the *J. albifrons* complex is not typical for the genus as a whole. Notwithstanding, the modest coverage of the *Jaera* species, our phylogenetic analysis reveals a wide discrepancy between morphological and molecular data for the genus. Our results suggest that the *J. albifrons* complex represents a relatively recent migration out of the Mediterranean basin and not an early-diverging lineage as suggested previously (*41*).

In the investigated populations, the proportion of hybrids determined morphologically generally did not exceed the values established for populations from other seas. In the Oleny Island population, however, the proportion of *albifrons*-*praehirsuta* hybrids was exceptionally high. In this population, the hybrid status of some males could not be determined with certainty, which is characteristic of the presence of backcrosses and introgression in the population (*10*, *49*). A small number of *albifrons*-*ischiosetosa* hybrids were also found here, but no *ischiosetosa*-*praehirsuta* hybrids. This asymmetry is particularly noteworthy in light of the fact that in this population, all three species inhabit without visible signs of stable spatial segregation.

Different factors are responsible for spatial separation of the species of the *J. albifrons* complex in the studied mixed settlements. Detailed analysis of the distribution of males in the Fucus Bay settlement on Ryazhkov Island revealed contrasting species ratios for two horizons within the fucoid belt, separated by only 12 m. However, at an intermediate level, the two species coexist in equal proportions. The difficulty of separating some *J. albifrons* specimens from *J. praehirsuta* using the rRNA genotyping data, combined with small sample sizes are likely responsible for the lack of power in the analysis of the spatial distribution of both males and females using genotyping data alone. For Yuzhnaya Bay, the factor of substrate was found crucial for segregation of *J. ischiosetosa* and *J. albifrons* with the former found on rocks and the latter significantly predominated on fucoids (see Table 20 and Fig. 29). In this case, this contrasting distribution of the species is evident in both the distribution of females carrying different genotypes on the two types of substrates and in a similar experimental design for genotyped males.

On Oleny Island, all three species cohabit a flat intertidal zone with a mixed composition of fucoids, but the fucoid species does not significantly influence the species composition. Instead, an unknown abiotic factor is responsible for changes in the proportion of *J. albifrons* and *J. ischiosetosa* males.

Thus, different factors influence the spatial segregation of individuals of different species in different mixed settlements. The influence of these factors can be partially discerned in the distribution of females carrying different genotypes. However, at least in some settlements, this segregation may be weak. This diversity precludes broad extrapolations from the studied settlements. However, the results obtained to some extent reflect the possible degrees of spatial segregation of species in the *J. albifrons* group.

## Acknowledgments

Most of the work was carried out in the Laboratory of Molecular Systematics of the Zoological Institute of the Russian Academy of Sciences and in the Kandalaksha State Nature Reserve: many thanks to Dr. Natalia Abramson (Zoological Institute) and Alexander Koryakin (Kandalaksha State Nature Reserve) who made this work possible. The authors express deep gratitude to everyone who participated in collecting the material for this work: Ksenia Shunkina, Maria Skazina, Sergey Korsun, Petr Strelkov, Vladimir Krapivin, Anton Vagapov (St. Petersburg State University), and Mikhail Fokin (Zoological Institute). Special thanks to Alexey Kostygov and Mikhail Fokin (Zoological Institute) for valuable discussions and assistance in the lab work. The work was supported within the framework of the Fundamental Research Program of the Presidium of the Russian Academy of Sciences “Biodiversity and Gene Pool Dynamics”.

