## Supplementary Material for "On rRNA gene cluster variation, phylogeny and local ecological differentiation in the *Jaera albifrons* complex (Crustacea: Isopoda)"

Andrey Rozenberg, Vadim Khaitov

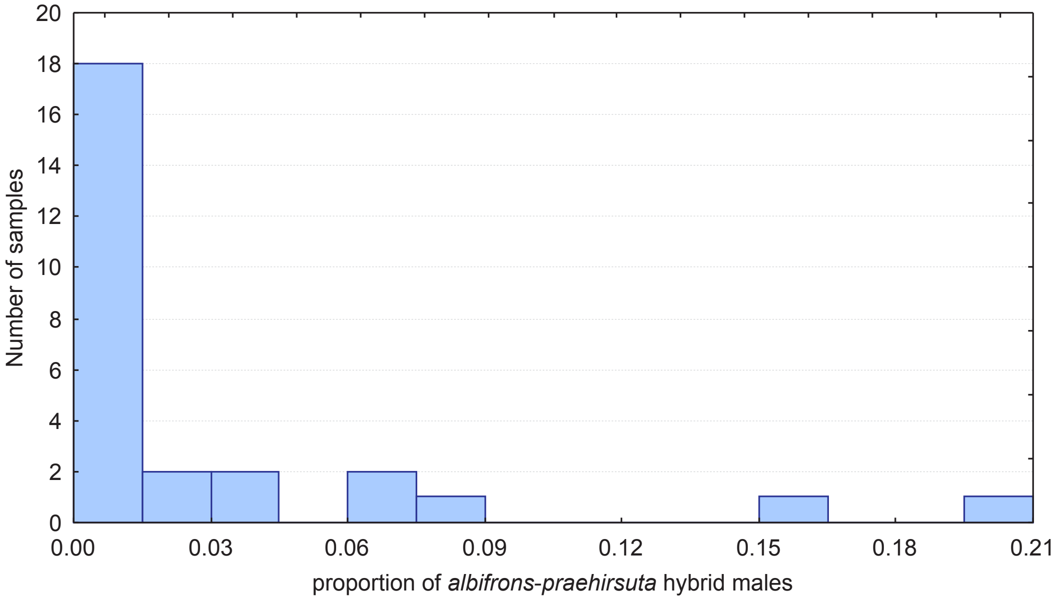


*Suppl. Figure 1*. Frequency distribution of samples (individual fucoid thalli) with different proportions of hybrid *albifrons-praehirsuta* males in the mixed settlement from Oleny Island.

*Suppl. Table 1*. Material used for genotyping of individuals of the *Jaera albifrons* complex: sample sizes are indicated for males of each species: A – *J. albifrons*, I – *J. ischiosetosa*; P – *J. praehirsuta*, hybrid males (AP and AI), and females. Individuals for which the D1-D2 fragment (11 individuals) and ITS1 (8) were partially or completely sequenced are marked with the letters D and S, respectively. Samples for which the D1-D2 fragment was sequenced with universal primers are marked with the letter C.

| Sea | Location | Collection date | A | I | P | AI | AP | females |
|---|---|---|---|---|---|---|---|---|
| White Sea | Chupa Bay | aquarium | | 1 | 2[DS] | | | |
| | Yuzhnaya Bay, Ryazhkov Island | 2008-06 | 22 | 35 | | 2 | | 41 |
| | Fucus Bay, Ryazhkov Island | 2008-06 | 40 | 1 | 30 | | 1 | 55 |
| | Dolgaya Bay, Bolshoy Solovetsky Island | 2009-07 | | | 2 | | | 2 |
| | Philip's Ponds and environs, Bolshoy Solovetsky Island | 2009-06 | 10 | 9 | 9 | 3 | | 10 |
| | Luvenga | 2008 | 2[DDS] | 1[DS] | | | | |
| | Oleny Island | 2007-06 | 2[DDS] | 1[DS] | 2[DDCC] | | | |
| Barents Sea | Abram-mys, Kola Bay | 2008 | | | 2[DSC] | | | 1 |
| | Yarnyshnaya Bay | 2009-08 | 6 | | | | | 7 |
| | Mogilnoye Lake, Kildin Island | 2007-07 | | 1[DSC] | | | | 3[CC] |
| | Zelenetskaya Bay | 2009-08 | | | 3 | | | 5 |
| Baltic Sea | Tvärminne | 2009-06 | 2 | 1 | | | | 6 |
| | Koporye Bay, Nennisari | 2007-11 | | | 1[S] | | | |
| North Sea | Bremerhaven | 2010-05 | | | | | | 2 |
| | Total: | | 85 | 48 | 47 | 5 | 1 | 132 |

*Suppl. Table 2.* Material used for comparative analysis of the nucleotide sequences of D1-D2, ITS1 and the 3' fragment of SSU.

| Species | Location | Collection date |
|---|---|---|
| *Jaera italica* | Adriatic Sea, Velja Rijeka stream at Prčanj | 2008-09 |
| *Jaera cornuta* | Adriatic Sea, Budva Bay | 2008-09 |
| *Jaera hopeana* | Adriatic Sea, Bay of Kotor | 2008-09 |
| *Jaera sarsi* | River Rhine, Bonn | 2010-05 |
| *Jaera caspica* | Middle Caspian Sea, 25-151 m depth | 2008-09/10 |
| *Carpias algicola* | Marine tropical aquarium at the Department of Invertebrate Zoology SPbSU | aquarium |
| *Asellus aquaticus* | Freshwater pool, park Alexandrino, St. Petersburg | 2006-10 |

*Suppl. Table 3.* Primers for amplification, sequencing, and genotyping used in this work. * – primers labelled with Cy5. “Strand” indicates the orientation of the primer relative to the coding strand: F for forward and R for reverse primers. “Position” indicates the position of the first nucleotide of the primer relative to the 5’ or 3’ end of the corresponding *Drosophila melanogaster* gene. Asterisk indicates primers designed here.

| Primer | Sequence | Strand | Position, bp | Target fragment | Specificity | Ref. | Mate primers |
|---|---|---|---|---|---|---|---|
| 1800 | GATCCTTCCGCAGGTTCACCTACG | R | 28 from 3’ SSU | SSU | universal | (*50*) | 1350F, 1800 |
| 1350F | CTGAAACTTAAAGGAATTGACG | F | 780 from 3' SSU | SSU | Pancrustacea? | * | 1800 |
| 18a1 | CCTAYCTGGTTGATCCTGCCAGT | F | ~1 от 5’ SSU | SSU, ITS1 | Pancrustacea? | (*50*) | 5p8aR, 1800 |
| 5p8aR | TCGACTCACGAGCCAAGTGATCC | R | 35 from 5’ 5.8S | ITS1, SSU | Asellota, other Pancrustacea | * | JISF, 18a1 |
| LSU D1, D2 fw1 | AGCGGAGGAAAAGAAACTA | F | 56 from 5’ LSU | D1-D2 LSU | universal | (*25*) | LSU D1, D2 rev1 |
| D12FJ | AGGAGAAGCTCAGCGTGTAGC | F | ~134 from 5’ LSU | D1-D2 LSU | Asellota | * | D12RN |
| D12FN | TTAAGCATATCASTAAGCGGAG | F | 41 from 5’ LSU | D1-D2 LSU | Isopoda | * | D12RN |
| LSU D1, D2 rev1 | TACTAGAAGGTTCGATTAGTC | R | 1111 from 5’ LSU | D1-D2 LSU | universal | (*25*) | LSU D1, D2 fw1 |
| D12RN | AGTTCCATACGCACTTCAAACA | R | ~1000 from 5’ LSU | D1-D2 LSU | Isopoda | * | D12FJ |
| J1187F | TAGCCAAGTTGCTGTAA* | F | | hairpin H32 in ITS1 | *J. albifrons* group | * | J1187R |
| J1187R | AGATCGAAAGGTCAAAA | R | | hairpin H32 in ITS1 | *J. albifrons* group | * | J1187F |
| J643F | TAAGGCTTGATTGTTGAG* | F | | position 643 in D1-D2 | *J. albifrons* group | * | J643R |
| J643R | ATAGGTAATGAGCGAAGG | R | | position 643 in D1-D2 | *J. albifrons* group | * | J643F |
| J842F | TGAAGTGTATGGAAGTA | F | | hairpin H27 in ITS1 | *J. albifrons* group | * | J842R |
| J842R | AATATGGGTATGCCGTG* | R | | hairpin H27 in ITS1 | *J. albifrons* group | * | J842F |
| JD426F | GCAGAAAGCGTGGTCGA | F | | position 426 in D1-D2 | *J. albifrons* group | * | D12RN |
| JISF | CGTAACAAGGTTTTCGTAGGTG | F | 43 from 3’ SSU | ITS1 | universal | * | 5p8aR, JISR |
| JISR | GCTACACGCTGAGCTTCTCCTG | R | 127 from 5’ LSU | ITS1, 5.8S, ITS2 | Asellota | * | JISF |

*Suppl. Table 4*. Nucleotide sequences used for phylogenetic reconstructions. GenBank accessions are indicated. Asterisks indicate sequences obtained here.

| Species | 3' SSU | ITS1 | D1-D2 | Family |
|---|---|---|---|---|
| *Asellus aquaticus* | AF255701 | MG751072* | EF417057 | Asellidae |
| *Paramunnopsis* sp. 1 | EF682227 | | EF682318 | Munnopsididae |
| *Eurycope* sp. MB I31 | EF682254 | | EF682323 | Munnopsididae |
| *Ianiropsis epilittoralis* | EF682260 | | EF682305 | "Janiridae" |
| *Carpias algicola* | MG751077* | MG751073* | * | "Janiridae" |
| *Jaera hopeana* | MG751080* | MG751075* | MG751084* | "Janiridae" |
| *J. albifrons* group | ×3*<br>AF279609<br>AF279610 | MG751076* | ×6* | "Janiridae" |
| *J. italica* | MG751078* | | MG751086* | "Janiridae" |
| *J. cornuta* | MG751079* | | MG751083* | "Janiridae" |
| *J. sarsi* | MG751081* | | | "Janiridae" |
| *J. caspica* | * | MG751074* | MG751082* | "Janiridae" |

*Suppl. Table 5*. ITS1 sites with inter-individual variability based on sequencing results in the *J. albifrons* group. Alternative states for each site are indicated.

| | 20 | 163 | 324 | 414 | 842 | 1187 |
|---|---|---|---|---|---|---|
| Variant 1 | $T_5$ | **A**TAT**CG**GACTGA-AGATT**ATAA** | G | T | $AC_4$ | $A_4$ |
| Variant 2 | $T_4$ | **T**TA**A**--GACTGA**C**AGATT**GA**A**C** | A | C | $AC_5$ | $A_3$ |

*Suppl. Table 6*. Distribution of *Jaera albifrons* and *J. praehirsuta* males in the Fucus Bay settlement. Results of ANOVA for the factors "level" and "site" with the estimated population densities (left) and species proportion (right) as dependent variables. Cochran's test for the significant factor: $p$ = 1.00. Here and in other tables, factor effects are indicated in parentheses: "R" for random and "F" for fixed. If not indicated otherwise, effects with significant contribution are highlighted in red ($p \leq 0.05$).

| Factor (effect) | *df* | Population density | | | | | | Species proportion | | |
|---|---|---|---|---|---|---|---|---|---|---|
| | | *J. albifrons* | | | *J. praehirsuta* | | | | | |
| | | *SS* | *F* | *p* | *SS* | *F* | *p* | *SS* | *F* | *p* |
| Site (R) | 1 | 0 | 0.34 | 0.618 | 0 | 0.14 | 0.748 | 0.15 | 3.3 | 0.211 |
| Level (F) | 2 | 0.004 | 1.9 | 0.345 | 0.008 | 12.12 | 0.076 | 2.46 | 27.32 | 0.035 |
| Interaction | 2 | 0.002 | 3.14 | 0.08 | 0.001 | 0.83 | 0.46 | 0.09 | 2.48 | 0.125 |
| Residual | 12 | 0.004 | | | 0.005 | | | 0.22 | | |

*Suppl. Table 7*. Proportions of males of *J. albifrons* and *J. praehirsuta* in the Fucus Bay settlement: Tukey's post-hoc test for the factor "Level". Levels in the intertidal zone are shown from upper to lower. Significant differences in means are highlighted in red ($p < 0.01$).

| Level | A | C | D | Average proportion of *J. albifrons* males |
|---|---|---|---|---|
| A | - | 0.0002 | 0.0002 | 1 |
| C | 0.0002 | - | 0.0016 | 0.462 |
| D | 0.0002 | 0.0016 | - | 0.11 |

*Suppl. Table 8*. Distribution of *J. albifrons* and *J. ischiosetosa* males on fucoids in the settlement from Yuzhnaya Bay. Results of ANOVA for the factors “level” (distance from water’s edge at low tide) and “stream” (stream bed/next to the stream) with population densities (left) and species proportion (right) as dependent variables.

| Factor (effect) | *df* | Population density | | | | | | Species proportion | | |
|---|---|---|---|---|---|---|---|---|---|---|
| | | *J. albifrons* | | | *J. ischiosetosa* | | | | | |
| | | *SS* | *F* | *p* | *SS* | *F* | *p* | *SS* | *F* | *p* |
| Stream (R) | 2 | 0.01 | 0.65 | 0.607 | 0.002 | 4.12 | 0.195 | 0.077 | 4.41 | 0.185 |
| Level (F) | 1 | 0.005 | 0.62 | 0.512 | 0 | 0.12 | 0.764 | 0 | 0.03 | 0.877 |
| Interaction | 2 | 0.016 | 0.7 | 0.515 | 0.001 | 0.71 | 0.512 | 0.017 | 0.63 | 0.551 |
| Residual | 12 | 0.135 | | | 0.004 | | | 0.167 | | |

*Suppl. Table 9*. Distribution of *J. albifrons* and *J. ischiosetosa* males in the settlement from Yuzhnaya Bay: results of the analysis of variance for the factors “site” and “substrate” (stream samples). The variable (proportion of *J. albifrons* males) was *arcsin*-transformed ($arcsin\sqrt{r}$). Cochran’s test for the significant factor: $p = 0.180$.

| Factor (effect) | *df* | *SS* | *F* | *p* |
|---|---|---|---|---|
| Site (R) | 2 | 0.014 | 0.19 | 0.838 |
| Substrate (F) | 1 | 6.02 | 170.24 | 0.006 |
| Interaction | 2 | 0.07 | 1.43 | 0.277 |
| Residual | 12 | 0.23 | | |

*Suppl. Table 10*. Distribution of *J. albifrons*, *J. ischiosetosa* and *J. praehirsuta* males in the Oleny Island settlement: results of ANOVA for the factors "site" and "fucoid species" with population densities (left) and species proportions (right) as dependent variables. Cochran's test for the significant effects: $p$ = 0.674 and 0.730, respectively.

| Factor (effect) | *df* | Population density | | | | | | | | | Species proportion | | | | | | | | |
|---|---|---|---|---|---|---|---|---|---|---|---|---|---|---|---|---|---|---|---|
| | | *J. albifrons* | | | *J. ischiosetosa* | | | *J. praehirsuta* | | | *J. albifrons* | | | *J. ischiosetosa* | | | *J. praehirsuta* | | |
| | | *SS* | *F* | *p* | *SS* | *F* | *p* | *SS* | *F* | *p* | *SS* | *F* | *p* | *SS* | *F* | *p* | *SS* | *F* | *p* |
| Site (R) | 2 | 1.82 | 5.63 | 0.069 | 1.31 | 4.01 | 0.111 | 0.02 | 0.62 | 0.581 | 0.94 | 103.8 | 0 | 1.09 | 33.5 | 0.003 | 0 | 0.03 | 0.967 |
| Fucoid (F) | 2 | 1.38 | 4.28 | 0.102 | 0.58 | 1.78 | 0.281 | 0.01 | 0.18 | 0.844 | 0.02 | 2.79 | 0.174 | 0 | 0.13 | 0.878 | 0.02 | 1.85 | 0.27 |
| Interaction | 4 | 0.64 | 0.16 | 0.385 | 0.66 | 1.15 | 0.366 | 0.06 | 1.08 | 0.394 | 0.02 | 0.45 | 0.771 | 0.06 | 1.67 | 0.2 | 0.02 | 1.04 | 0.413 |
| Residual | 18 | 2.63 | | | 2.57 | | | 0.26 | | | 0.18 | | | 0.17 | | | 0.11 | | |

*Suppl. Table 11*. Proportions of male *J. albifrons* and *J. ischiosetosa* at different sites in the Oleny Island settlement: Tukey's post-hoc test. Significant differences in means ($p \leq 0.001$) are highlighted in red.

| Site | Proportion of *J. albifrons* | | | | Proportion of *J. ischiosetosa* | | | |
|---|---|---|---|---|---|---|---|---|
| | 1 | 2 | 3 | Average per site | 1 | 2 | 3 | Average per site |
| 1 | - | 0 | 0 | 0.084 | - | 0 | 0 | 0.822 |
| 2 | 0 | - | 0.987 | 0.475 | 0 | - | 0.882 | 0.385 |
| 3 | 0 | 0.987 | - | 0.482 | 0 | 0.882 | - | 0.407 |

*Suppl. Table 12*. Proportions of males of three species in the mixed settlement from Oleny Island: results of PERMANOVA analysis of variance for three variables representing the proportions of males of each species. Proportions were *arcsin*-transformed (*arcsin*√*r*). Multivariate analogue of Levene's homoscedasticity test (PERMDISP) for the significant effect: $p = 0.068$.

| Factor (effect) | *df* | *SS* | *F* | *p* |
|---|---|---|---|---|
| Site (R) | 2 | 2.68 | 26.35 | 0.0002 |
| Fucoid species (F) | 2 | 0.11 | 0.93 | 0.606 |
| Interaction | 4 | 0.23 | 1.14 | 0.362 |
| Residual | 18 | 0.91 | | |

*Suppl. Table 13.* The success of posteriori species identification for males based on quantitative rRNA genotypes using discriminant analysis.

| Settlement | Identifications | *albifrons* | *ischiosetosa* | *praehirsuta* |
|---|---|---|---|---|
| B. Solovetsky Island | total | 10 | 9 | 9 |
| | incorrect | 1 | 0 | 0 |
| Ryazhkov Island | total | 61 | 37 | 30 |
| | incorrect | 7 | 0 | 7 |

*Suppl. Table 14*. Spatial distribution of females in the two Ryazhkov Island settlements based on genotyping results: PERMANOVA results for the factors “site” and “level”. Tests for a similar sample of males are provided for comparison. For Yuzhnaya Bay, values were ∜-transformed. Permutation values are shown as significance levels for all factors except for “substrate” in Yuzhnaya Bay, for which *p*-values were obtained with the Monte Carlo method. Multivariate analogue of Levene’s homoscedasticity test (PERMDISP) for the significant factors: $p = 0.185$ (Fucus Bay, interaction term, females), 0.808 (Yuzhnaya Bay, “substrate”, females) and 0.020 (Yuzhnaya Bay, “substrate”, males).

| Factor | *df* | females | | | males | | |
|---|---|---|---|---|---|---|---|
| | | *SS* | *F* | *p* | *SS* | *F* | *p* |
| Fucus Bay | | | | | | | |
| Site (R) | 1 | 0.008 | 0.391 | 0.816 | 0.036 | 2.009 | 0.113 |
| Level (F) | 2 | 0.048 | 0.51 | 0.821 | 0.052 | 0.876 | 0.531 |
| Interaction | 2 | 0.094 | 2.11 | 0.034 | 0.059 | 1.674 | 0.128 |
| Residual | 36 | 0.769 | | | 0.637 | | |
| Yuzhnaya Bay | | | | | | | |
| Site (R) | 1 | 0.12 | 0.41 | 0.592 | 0.05 | 0.67 | 0.67 |
| Substrate (F) | 1 | 3.82 | 80.7 | 0.004 | 4.38 | 15.12 | 0 |
| Interaction | 1 | 0.05 | 0.17 | 0.822 | 0.07 | 0.25 | 0.734 |
| Residual | 24 | 6.75 | | | 6.93 | | |

*Suppl. Table 14*. Spatial distribution of females in the Fucus Bay settlement based on genotyping results: post-hoc *t*-test for the interaction between “level” and “site”. *p*-values are shown for differences between different levels at the two sites. Significant differences are highlighted in red ($p \leq 0.017$, taking into account Bonferroni correction).

| Site | Level | A | C | D |
|---|---|---|---|---|
| 1 | A | - | 0.359 | 0.906 |
| | C | 0.359 | - | 0.762 |
| | D | 0.906 | 0.762 | - |
| 2 | A | - | 0.029 | 0.008 |
| | C | 0.029 | - | 0.184 |
| | D | 0.008 | 0.184 | - |
